# Batch Effect Correction of RNA-seq Data through Sample Distance Matrix Adjustment

**DOI:** 10.1101/669739

**Authors:** Teng Fei, Tianwei Yu

## Abstract

Batch effect is a frequent challenge in deep sequencing data analysis that can lead to misleading conclusions. We present scBatch, a numerical algorithm that conducts batch effect correction on the count matrix of RNA sequencing (RNA-seq) data. Different from traditional methods, scBatch starts with establishing an ideal correction of the sample distance matrix that effectively reflect the underlying biological subgroups, without considering the actual correction of the raw count matrix itself. It then seeks an optimal linear transformation of the count matrix to approximate the established sample pattern. The benefit of such an approach is the final result is not restricted by assumptions on the mechanism of the batch effect. As a result, the method yields good clustering and gene differential expression (DE) results. We compared the new method, scBatch, with leading batch effect removal methods ComBat and mnnCorrect on simulated data, real bulk RNA-seq data, and real single-cell RNA-seq data. The comparisons demonstrated that scBatch achieved better sample clustering and DE gene detection results.

## Introduction

In the recent decade, RNA sequencing (RNA-seq) has become a major tool for transcriptomics. Due to the limitation of sequencing technology and sample preparations, technical variations exist among reads from different batches of experiments. These unwanted technical variations, or batch effects, can lead to misleading scientific findings in downstream data analysis (Hicks et al., 2017). Typically, batch effects can alter the sample patterns, causing false interpretations about cell lineage and heterogeneity. If the goal is to detect differential expression (DE) genes, the analysis can suffer loss of statistical power and/or bias.

While the severity of batch effects varies in different datasets, batch effect corrections were shown to be effective in general. For instance, batch effect correction on the ENCODE human and mouse tissues bulk RNA-seq data (Lin et al., 2014), where the batch effects were intense, obtained largely different and more sensible tissue clustering results compared to before correction (Gilad and Mizrahi-Man, 2015). In other datasets, batch effects are often more subtle. In such cases, although the true biological pattern is maintained to some extent, weak to moderate batch effects can still be observed. Hicks et al. (2017) discussed the coexistence of biological signal and technical variation, which may still compromise the downstream analysis. The correction of the batch effects can yield better clustering results (Fei et al., 2018) on data with weak to moderate batch effects that were unobvious from dimension reduction plots (Usoskin et al., 2015; Muraro et al., 2016). These previous efforts argue for the inclusion of batch effect corrections as a routine procedure in data preparation.

Since the microarray era, efforts have been made to correct batch effects. Johnson et al. (2007) proposed an empirical Bayes algorithm, ComBat, to normalize the data by removing additive and multiplicative batch effects, which continued to be a successful method in RNA-seq data. Researchers also attempted to find and correct unknown batch effects by utilizing control genes in microarray (Gagnon-Bartsch and Speed, 2012) and RNA-seq data (Leek, 2014; Risso et al., 2014; Chen and Zhou, 2017). ComBat and the control gene methods are based on regression models, while more recent methods proposed different strategies that allow for more complex batch effect mechanisms. To achieve better clustering performance, Fei et al. (2018) developed a non-parametric approach, named QuantNorm, to correct sample distance matrix by quantile normalization; Haghverdi et al. (2018) utilized the mutual nearest neighbor relationships among samples from different batches to establish the MNN correction scheme. As observed from the results in Fei et al. (2018) and Haghverdi et al. (2018), current methods have reached reasonable performances in sample pattern detection, such as finding clusters or conducting dimension reduction.

However, DE detection appears not to be the emphasis of recent methods development. Chen and Zhou (2017) only evaluated the clustering performances, while QuantNorm (Fei et al., 2018) only returns corrected distance matrices and does not support DE tests. Although Haghverdi et al. (2018) conducted DE tests, the user manual (https://bioconductor.org/packages/3.8/workflows/vignettes/simpleSingleCell/inst/doc/work-5-mnn.html) of the corresponding bioconductor function, mnnCorrect, recommends not using the corrected count matrix for DE analysis with considerations on manipulated data scales and mean-variance relationship.

Motivated by the challenges faced in DE detection, in this study we develop a new method, sc-Batch, to utilize the corrected sample distance matrix to further correct the count matrix. Specifically, we seek a linear transformation to the count matrix, such that the Pearson correlation matrix of the transformed matrix approximates the corrected correlation matrix obtained from QuantNorm. For this purpose, we propose a random block coordinate descent algorithm to conduct linear transformation on the *p* (genes) × *n* (samples) count matrix. Simulation studies demonstrate that in terms of DE gene detection, our method corrects the count matrix better compared to ComBat and MNN, with consistently higher area under the receiver operating characteristic curve (AUC) and area under the precision-recall curve (PRAUC). In real data analyses, the proposed method also show strong performances in clustering and DE detection in a bulk RNA-seq dataset (Lin et al., 2014) and two scRNA-seq datasets (Usoskin et al., 2015; Xin et al., 2016).

## Results

### Batch effect correction based on corrected sample correlation matrix

The scBatch method considers a study design scenario where the cell type or disease status composition is not severely confounded with batch, i.e. different cell subtypes or disease status are roughly evenly distributed among the batches. This balanced study design scheme has been recommended (Hicks et al., 2017) and widely adopted because it helps to avoid bias caused by confounding with batch. Under this assumption, although the batch effect may interrupt the overall data pattern, the data pattern within each batch should share similar characteristics, including similar quantile distributions in different batch blocks in the sample distance matrix. Moreover, the balanced study design retain the above feature in various types of omics data, allowing our approach to be applied to different types of data without restrictions of distribution assumptions on the distance matrix.

Fig. 1 summarizes the workflow of scBatch. Given a count matrix *X* and its Pearson correlation matrix, we first utilize QuantNorm to obtain the corrected sample Pearson correlation matrix *D*. Then *X* and *D* are input to the proposed algorithm to seek the weight matrix *W*, such that the Pearson correlation matrix of the linear transformation *X* × *W* approximates *D*. After the algorithm converges, the linear-transformed count matrix *Y* = *X* × *W* is output as the corrected count matrix that inherits the corrected sample pattern in *D*. Although more complex models can be used to achieve nonlinear transformation, we believe linear transformation can avoid over-correction while still achieving good results. Detailed problem setup and algorithm design can be found in the Methods section.

**Figure 1:**
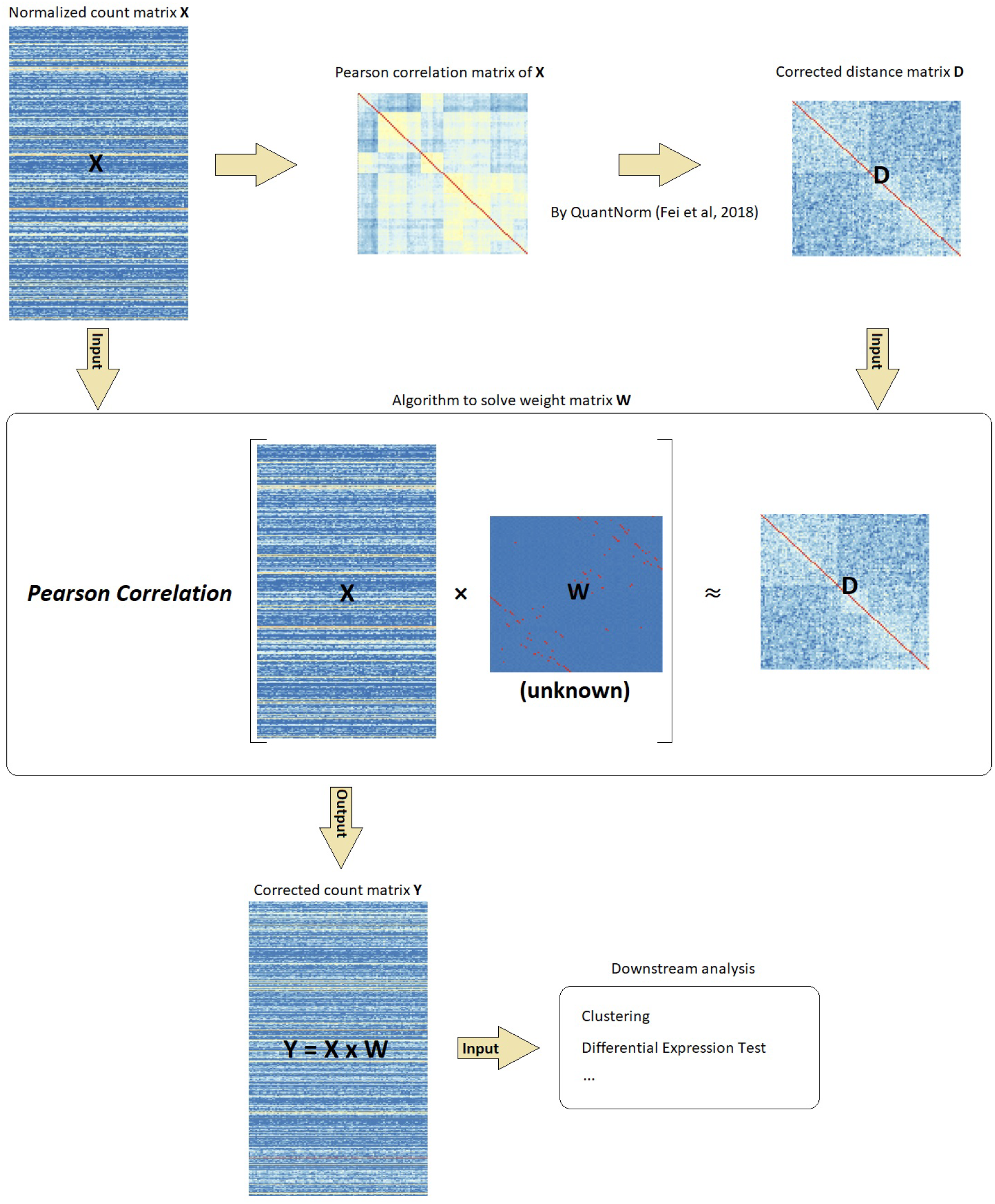
Overview of scBatch workflow. For the preprocessed count matrix *X*, the Pearson correlation matrix is corrected by QuantNorm to obtain a reference sample distance matrix *D*. Then the main algorithm is utilized to search for the weight matrix *W* to achieve the objective that the Pearson correlation of *X* × *W* is close to *D*. The corrected count matrix *Y* = *X* × *W* inherits the sample pattern information from *D*, which can be used for downstream analyses.

### scBatch achieves better DE detection in simulations

In previous studies, we have shown that the corrected distance matrix by QuantNorm, i.e. the reference matrix used in this study, achieved better sample clustering results in simulations (Fei et al., 2018). However QuantNorm could not correct the count matrix, and hence the inability to improve differentially expressed (DE) gene detection. In this simulation study, we focused on the capability of the new method to facilitate the detection of DE genes after count matrix correction.

Simulated scRNA-seq datasets were generated by Bioconductor package `splatter` (Zappia et al., 2017) which controls the batch effect mechanisms by customizing relevant parameters. For all simulated datasets, four types of cells were distributed by random sampling in 4 batches. To compare the effectiveness of DE detection using uncorrected data, MNN correction, ComBat correction and scBatch correction, we considered five configurations of batch effect with different mechanisms (Fig. 2). For configurations (I) to (IV), the complexity of batch effects increases, while configuration (V) serves as the control group where no batch effect was introduced. Detailed simulation design and data generation procedures are reported in Methods.

**Figure 2:**
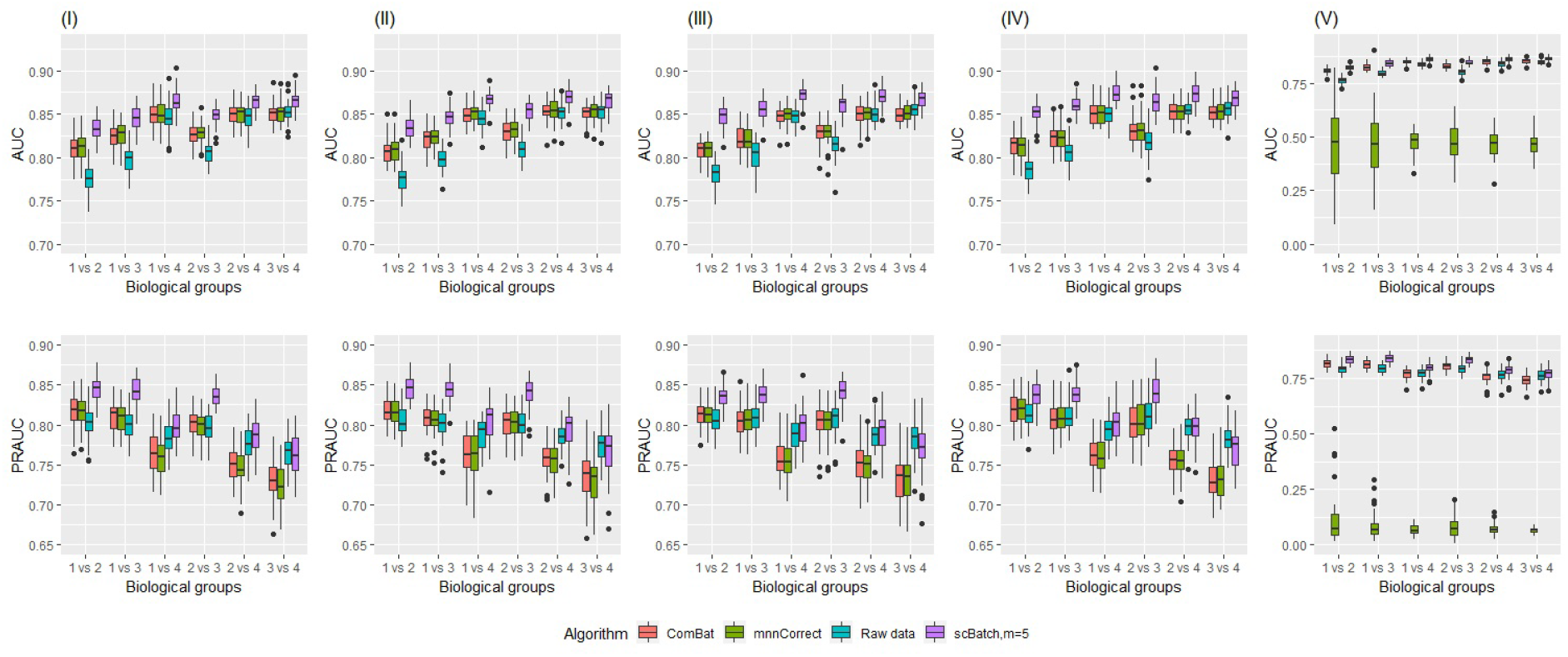
Boxplots of AUC and PR-AUC indices computed from adjusted p-values from DE tests in 50 simulations from the five configurations (I - V). Each column represents one corresponding configuration. The batch effect complexity was increased from configuration (I) to configuration (IV). Configuration (V) contained no batch effect.

Standard DE detection method Seurat (Satija et al., 2015) was utilized to conduct DE gene detection for each pair of the four cell types. The available gold standard of DE gene lists enabled us to compute area under the receiver operating characteristic curve (AUC) and area under the precision-recall curve (PR-AUC), based on the adjusted p-values from DE tests. At each parameter setting, the simulations and tests were repeated 50 times to obtain 50 corresponding AUC and PR-AUC values.

Fig. 2 displays the boxplots, generated by `ggplot2` (Wickham, 2016), of AUCs and PR-AUCs for pairwise DE tests obtained from the 50 simulations. While ComBat and MNN achieved some improvement from the uncorrected data, scBatch consistently ranked at the top in both metrics under different simulation settings. Interestingly, under the control configuration where there was no batch effect, MNN correction failed to retain the original sample pattern, resulting in reduced power in DE detection.

### scBatch obtained better sample patterns for bulk RNA-seq data

To illustrate the utility of scBatch on different types of real data, we first considered a bulk RNA-seq data of human and mouse tissues (Lin et al., 2014). The dataset consists of 13 tissues from human and their counterparts in mouse. It is a typical example of data containing strong batch effects. Without batch effect correction, the samples from the same species clustered together, as the tissues from different species were measured in different batches. Several re-analyses have shown that the samples would be clustered by tissues instead of species after proper batch effect correction (Gilad and Mizrahi-Man, 2015; Sudmant et al., 2015; Fei et al., 2018).

In our analysis, we compared the clustering performance of batch effect correction algorithms, including ComBat, MNN and scBatch. We used heatmaps generated by R package `pheatmap` (Kolde and Kolde, 2015) and Adjusted Rand Index (ARI) (Hubert and Arabie, 1985) calculated from hierarchical clustering for the comparison. As observed in Fig. 3A, the subjects were mainly clustered by species in the raw count matrix. After batch effect correction, ComBat (ARI = 0.70, Fig 3B) and MNN (ARI = 0.68, Fig 3C) restored 10 out of 13 pairs of tissues, while scBatch (ARI = 0.88, Fig 3D) out-perfomed the other two methods by retrieving 12 out of the 13 matches with reasonably high contrast in the heatmap. The above results demonstrated scBatch’s ability to obtain better sample patterns. For scBatch, the restoring of sample patterns was based on our previously published work of sample correlation matrix correction (Fei et al., 2018). Thus the results here on the mouse/human tissue data only demonstrated that scBatch could truly adjust the count matrix such that the resulting sample correlation matrix approaches that generated by Fei et al. (2018). In the next sections, we demonstrate the utility of scBatch in DE gene detection on single cell RNAseq data.

**Figure 3:**
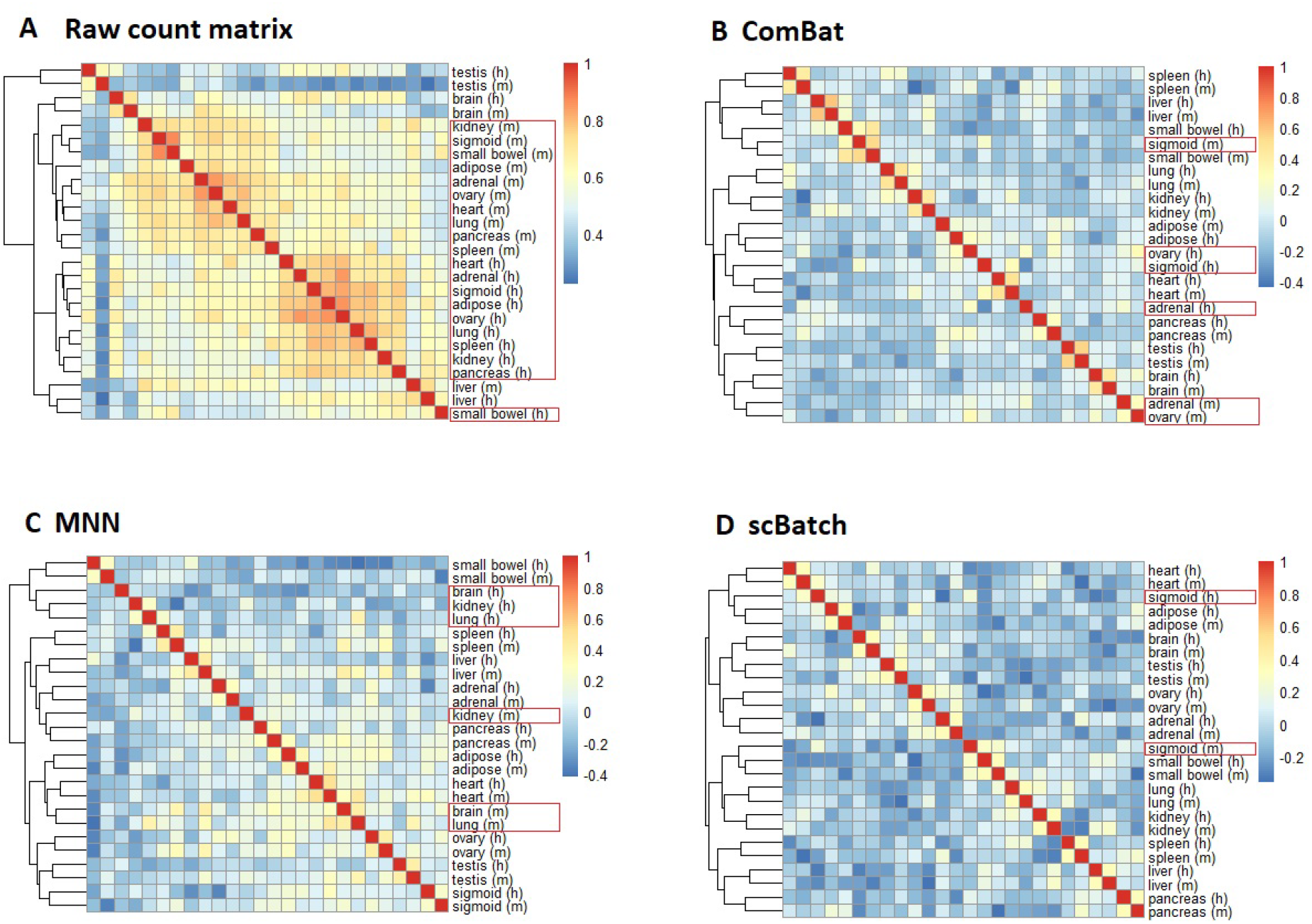
Heatmaps of ENCODE mouse and human tissues data generated from the Pearson correlation matrices of **A** raw count matrix; **B** count matrix corrected by ComBat; **C** count matrix corrected by MNN; **D** count matrix corrected by scBatch. Hierarchical clustering was used to generate clusters. Red rectangles mark the samples not clustered with their counterparts in the other species.

### scBatch shows strong performance in cell heterogeneity investigation

Under the assumption of scBatch algorithm, the correction can be reliably applied for batches with similar cell type compositions. This assumption is particularly suitable for the investigation of cell heterogeneity from a certain tissue. We compared clustering and DE gene selection results of different correction methods on two single-cell RNA-seq datasets, one for mouse neuron cells (Usoskin et al., 2015) and one for human pancreas cells (Xin et al., 2016). Our method not only obtained better sample patterns, but also retained important information in marker genes.

### Mouse neuron dataset GSE59739

The single-cell RNA-seq data was generated by Usoskin et al. (2015). Cell labels determined by marker genes were provided in the data. We based our analyses on the given cell labels to investigate the differences of four main subtypes of cells, namely non-peptidergic nociceptors (NP), tyrosine hydroxylase containing (TH), neurofilament containing (NF) and peptidergic nociceptors (PEP). As observed in the two-dimensional t-SNE plot (top-left panel in Figure 4A), the uncorrected data maintained the cluster of NF cells, while the other three subtypes formed a mixture. Regarding the library labels as batches, we conducted batch effect correction using ComBat, MNN and scBatch.

**Figure 4:**
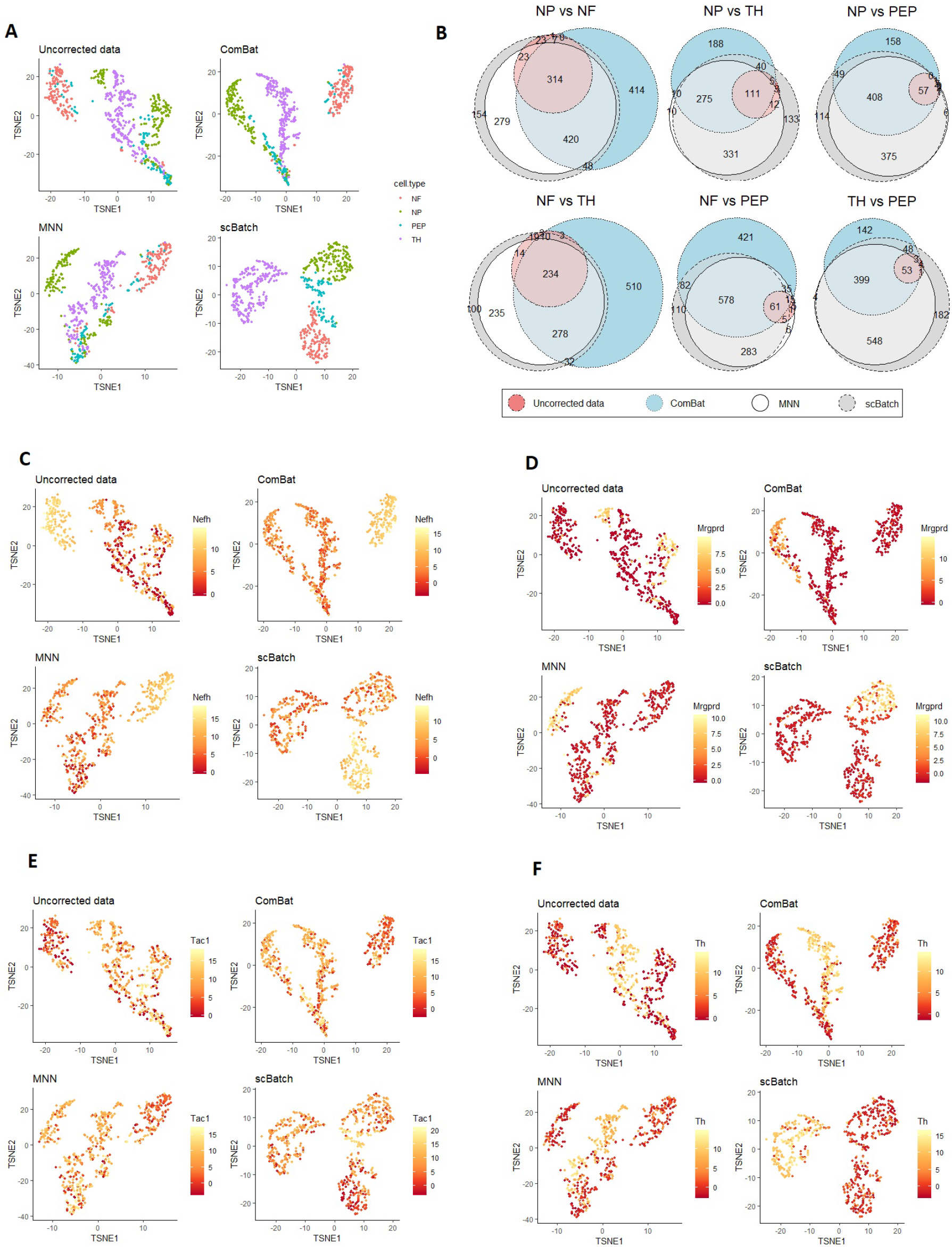
**A,C-F** t-SNE plots of the sample patterns from uncorrected data (normalized raw count data), ComBat correction, MNN correction and scBatch correction, colored by **A** cell labels from raw data, **C** marker gene *Nefh* for NF cells, **D** marker gene *Mrgprd* for NP cells, **E** marker gene *Tac1* for PEP cells, and **F** marker gene *Th* for TH cells. **B** Venn diagrams for the significant genes from pairwise differential gene tests by Seurat (Satija et al., 2015) with adjusted p-values < 10^*−*6^ and fold changes > 2.

In order to evaluate the clustering performance, we utilized t-distributed stochastic neighbor embedding (t-SNE) (Maaten and Hinton, 2008) dimension reduction, and the average ARI based on multiple k-means clustering results. As the t-SNE plots (Fig. 4A) display, scBatch (bottom right) obtained a clearer sample pattern which distinguished the four subtypes better. The ARI indices based on k-means clustering results also demonstrated that scBatch 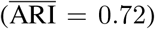 outperformed uncorrected data 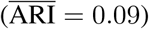, 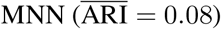 and 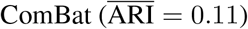 by a large margin. To further investigate whether corrected count matrices kept crucial marker information, we plot the marker gene expression levels in the t-SNE plots for the four cell subtypes, displayed in Figs. 4C, D, E, F. It can be observed that scBatch correction inherited the marker information from the uncorrected data with large contrast, while ComBat and MNN did not maintain as strong contrast in the marker genes.

Due to its ability to restore better sample patterns and maintain important marker contrasts, scBatch also showed good performance in DE gene detection. We conducted DE gene detection between all cell type pairs, using the method Seurat (Satija et al., 2015). Given the large differences between the cell types, we used stringent criteria of adjusted p-value < 10^*−*6^ and fold change > 2. Fig. 4B shows the Venn diagrams of DE genes detected from the different count matrices. As observed, the DE genes from the uncorrected data were largely contained in the DE gene set from the scBatch-corrected data. Moreover, scBatch found the largest number of DE genes in five out of the six subtype pairs, indicating that more underlying information masked by batch effects may be revealed by scBatch.

Between the three batch-effect correction methods, scBatch and MNN tended to agree with each other on this dataset. In most cases the majority of the genes found by MNN were also found by scBatch. ComBat disagreed with the other two methods in some of the comparisons. In order to examine their differences at the functional level, we used GOstats to analyze the over-representation of gene ontology biological processes by the selected genes (Falcon and Gentleman, 2007). In the neurofilament containing (NF) cells v.s. tyrosine hydroxylase containing (TH) cells comparison, the largest difference between ComBat and the other two methods was observed (Fig.4B). Functional analysis revealed that the top biological processes found in uncorrected data fall into axonogenesis, cell morphogenesis and synapse assembly (Supplemental File S1). scBatch and MNN identified similar top biological processes such as those involved in synapse assembly and cell adhesion. These results are reasonable given the functionality of the NF cells. On the other hand, the top terms resulting from the application of ComBat were focused on regulation of ion transmembrane transport and synapse organization, the first of which was not obvious in terms of the biological functions to the NF and TH cells. The full GOstats results are contained in Supplemental File S1.

### Human pancreas data GSE81608

We analyzed another single-cell RNA-seq data of human pancreas cells (Xin et al., 2016). The dataset consists of cells from healthy controls and patients with type II diabetes. In this study, cells from different donors were separately processed (Xin et al., 2016). Donor IDs can be regarded as batch labels. Here we focused on healthy control cells to investigate the cell heterogeneity. There are four dominating endocrine cell types - alpha cells that produce glucagon, beta cells that produce insulin and amylin, delta cells that produce somatostatin, and gamma cells that produce pancreatic polypeptide. In this dataset, the distribution of cell types between the batches vary substantially. The proportion of alpha cells in each batch ranged from 16.7% to 74.6% among the batches. The proportion of beta cells ranged from 14.0% to 54.2%. Given that gamma and delta cells account for small proportions in the pancreas islet, the were not present in some of the batches. The range for delta cell was 0 to 8.3%, and the range for gamma cell was 0 to 20.0%. These variations made the data more challenging than the mouse neuron data. As observed in Fig. 5A, the uncorrected data (top left) showed high distinction of alpha and beta cells, while the gamma and delta cells were clustered together.

**Figure 5:**
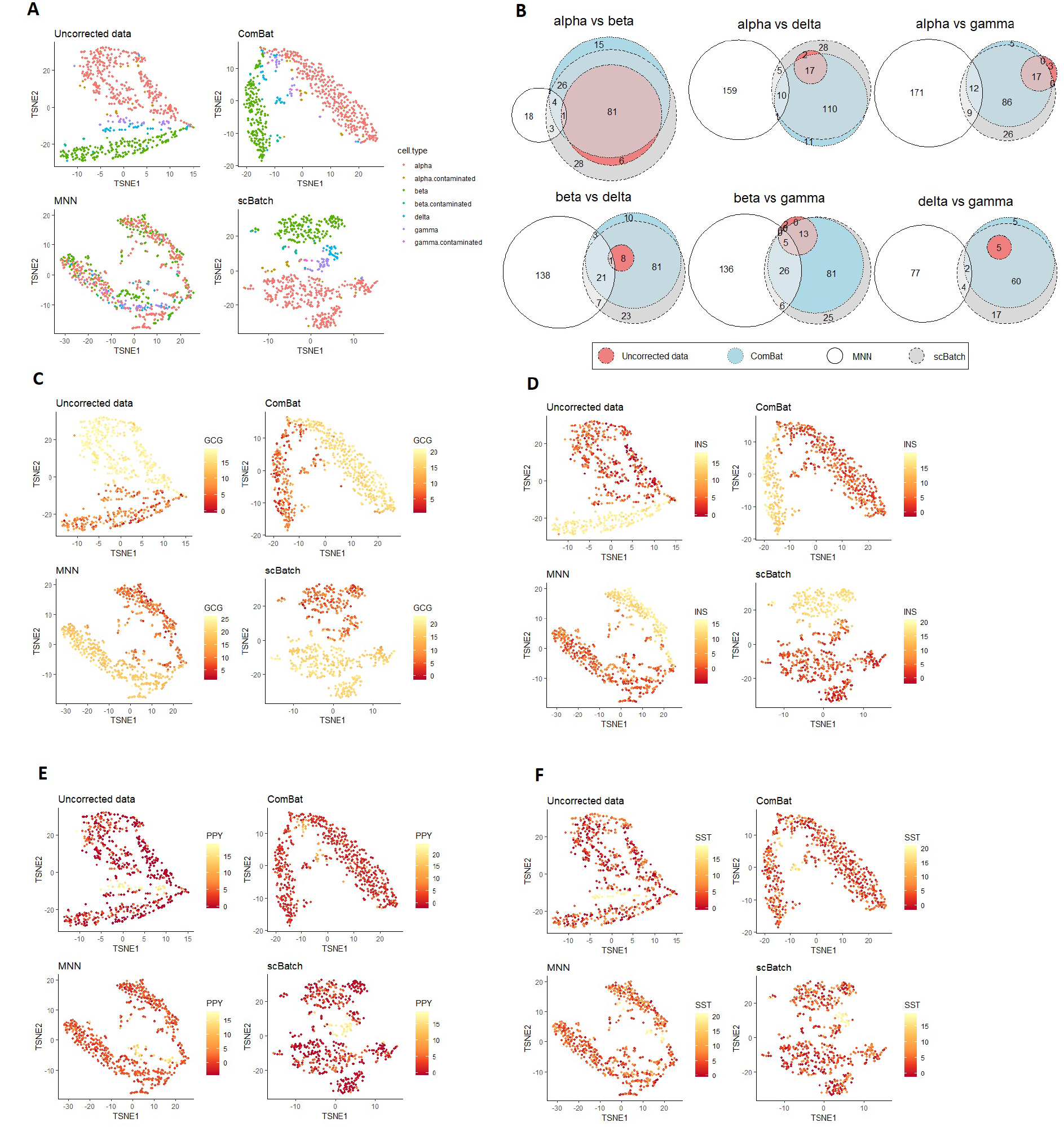
**A,C-F** t-SNE plots of the sample patterns from uncorrected data (normalized raw count data), ComBat-corrected data, MNN-corrected data and scBatch-corrected data, colored by **A**, cell types, **C**, marker gene *GCG* for alpha cells, **D**, marker gene *INS* for beta cells, **E**, marker gene *PPY* for gamma cells, and **F**, marker gene *SST* for delta cells. **B**, Venn diagrams for the significant genes from pairwise differential gene tests by Seurat (Satija et al., 2015) with adjusted p-values < 10^*−*6^ and log fold changes > 2.

We applied similar analysis procedure used for the mouse neuron data. From the t-SNE plots where cells were colored by the provided subtype labels (Fig. 5A), mixed gamma and delta cells appeared in the patterns for both uncorrected data (top left) and ComBat-corrected data (top right), while MNN-corrected data (bottom left) formed clusters which contained mixed types of cells. The pattern obtained by scBatch (bottom right), in contrast, successfully separated the four types of cells, although the distance between gamma and delta cells were still close.

As expected, in K-means clustering results, scBatch achieved highest average ARI (0.60), compared to uncorrected data (0.42) and ComBat (0.44). On the other hand, the average ARI for MNN (−0.01) indicates no correlation between the MNN sample pattern and the cell subtype labels. The marker gene expressions on t-SNE plots (Figs. 5C, D, E, F) similarly demonstrated that scBatch was able to maintain marker gene information from the original data. The imbalance between the batches, especially the lack of some cell types in certain batches, is likely the reason for MNN’s inconsistent performance. Its nearest-neighbor based approach may mistakenly match different type of cells, and cause the adjustment to be erraneous. On the other hand, although scBatch prefers balanced design, it appeared to be more robust against the imbalance.

In the Venn diagrams of significant DE genes (Fig. 5B), as the cell labels did not match the clusters in the MNN pattern, the detected DE gene set for MNN was largely different from the other three approaches. Apart from the results by MNN, high agreements were observed among uncorrected data, ComBat and scBatch, where scBatch was able to detect more DE genes in all six pairs.

We again used GOstats to analyze the over-representation of gene ontology biological processes by the selected genes (Falcon and Gentleman, 2007). We take the DE genes between alpha and beta cells as an example. The two types of cells produce hormones of opposite effects - glucagon and isulin, respectively. From uncorrected data, the top GO terms represented by the DE genes include “regulation of system process”, “digestion”, and “regulation of heart contraction”. From ComBat-corrected data, the top GO terms include “response to biotic stimulus”, “digestion”, and “regulation of system process”. The top GO terms from scBatch include “G-protein coupled receptor signaling pathway”, “response to lipopolysaccharide”, “digestion”. Lipopolysaccharides can modulate glucose metabolism (Nguyen et al., 2014). It is well-known that G-protein coupled receptors are regulated in beta cells to affect insulin secretion and their natural ligands (Persaud, 2017). The top GO terms from MNN were instead focused on immune-related processes, including “chemokine-mediated signaling pathway”, “inflammatory response” and “neutrophil chemotaxis”. They do not reflect the biological functions and differences of the alpha and beta cells. These results clearly indicate the scBatch and ComBat results are more biologically plausible. The full list of the functional analysis results are in Supplemental File S2.

## Discussion

Batch effects are frequently encountered in omics data analysis, thus a crucial issue to address before downstream analysis that leads to scientific discoveries. In this paper, we introduced a novel method for batch effect correction.We have shown that the proposed method, scBatch, can obtain better clustering pattern, maintain crucial marker information and detect more DE genes.

The method assumes roughly balanced sample population among batches. The assumption is strong yet reasonable (Hicks et al., 2017), and the method appeared to be robust when the assumption is mildly violated. In the presence of technical variations and various confounders, a randomized design is ideal. Data analysis requires proper randomization to determine the existence of latent groups or cell subtypes, or to decide whether the gene expression differs between two populations. In reality, data analysts sometimes encounter data where the biological groups and batch labels are totally confounded, which brings tremendous challenges in downstream analysis. On the other hand, if the study design is balanced, we will be able to restore the biological pattern, even if the technical variations dominate the sample pattern in the raw data.

Computational cost is another practical concern for the application of algorithms. Compared to ComBat and MNN, scBatch requires more computation time to reach optimal results. As shown in Supplemental Fig.1, the running time increases faster than a linear growth rate as the sample size increases. Moreover, for a fixed sample size, the time to convergence for scBatch also varied. The varied running time was determined by the complication of batch effects, which decided the similarity between uncorrected sample pattern and the corrected referencing sample pattern. Although slower than ComBat and MNN, the running speed of scBatch is within acceptable range. For a few hundred cells, the computing time was in the range of minutes. For large studies with over 1000 cells, the computing can take hours.

There is still large room to improve the proposed method. First, we only adopted the simplest linear transformation of raw count matrix in this paper, while a non-linear transformation may better depict the sample pattern in the corrected distance matrix. Secondly, the metrics of distance can also affect the correction. We used the Pearson correlation matrix because it was easy to interpret and convenient for gradient computation, while other distance metrics such as Spearman correlation may bring other insights to the data pattern. Thus, a more universal numerical gradient descent algorithm may be applied to adapt to different types of distance matrices.

## Methods

### Main algorithm

#### Problem setup

The count matrix *X*_*p*×*n*_ with *p* genes and *n* cells, in which the *n* cells fall into multiple batches, is subject to batch effects. Based on the Pearson correlation matrix of *X*, a corrected *n* × *n* correlation matrix *D* with improved sample similarities is obtained using the distance matrix correction algorithm QuantNorm (Fei et al., 2018). Given *D*, the objective is to solve for an optimal *n* × *n* weight matrix *W* such that the Pearson correlation of the linear-transformed count matrix *Y* = *XW* approximates the sample pattern in *D*. The transformed count matrix *Y* can then be used in downstream analyses. We note that similar to other methods, the resulting matrix may no longer be composed of non-negative integers.

#### Least squares loss function

In order to solve *W*, we propose to minimize the following least squares loss function

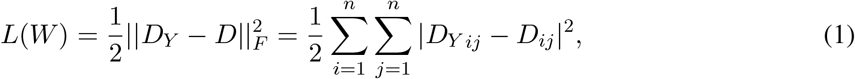

where *D*_*Y*_ is the Pearson correlation matrix of *Y*, ||⋅||_*F*_ is the Frobenius norm and *A*_*ij*_ denotes the (*i*, *j*) entry of matrix *A*. Thus, the optimized weight matrix *W*_*opt*_ satisfies *W*_*opt*_ = argmin_*W*_*L*(*W*) and the corrected count matrix is *Y*_*opt*_ = *XW*_*opt*_.

#### Gradient of the loss function

By chain rule, the gradient of the loss function *L*(*W*) is

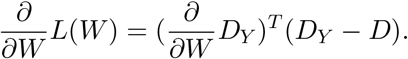

By definition, the *i*, *j* entry of *D*_*Y*_ satisfies

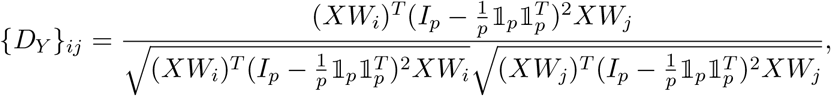

where *W*_*i*_ is the *i*th column of *W*, *I*_*p*_ is the *p* × *p* identity matrix, and 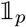 is the *p* × 1 vector with all entries equal to one. As can be observed, 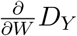 is a 4th-rank tensor in *n*-dimensional space. Thus the gradient of the loss function *L*(*Y*), which is the product of 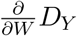 and the *n* × *n* matrix (*D*_*Y*_ − *D*), is also a *n* × *n* matrix.

Although the scale of computation appears large, we derived an equivalent but more economic approach to compute the gradient in practice. Since {*D*_*Y*_}_*ij*_ involves only two columns from *W*, the tensor 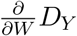 is sparse so that all its entries can be saved in a 3rd-rank tensor in *n*-dimensional space. Let *A*_*k*_ denote the *k*th column of matrix *A*. Considering the gradient performance and practical computing, moreover, we further decompose the calculation into columnwise gradients 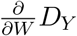, *k* = 1, …, *n*, which are *n* × *n* matrices. Using columnwise gradients as the unit, both coordinate gradient descent (Wright, 2015) and standard gradient descent can be easily implemented.

Denote 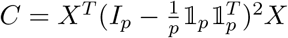. By some algebra, the columnwise gradient 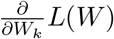 satisfies

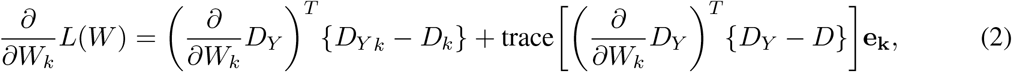

where **e**_*k*_ is a *n* × 1 vector in which the *k* entry is equal to one and others are equal to zero, and

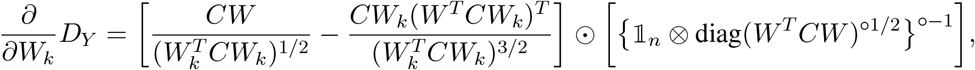

where ⊙, ∘ respectively represents Hadamard (elementwise) product and power, and ⊗ represents outer product.

##### Algorithm 1 Random block coordinate descent algorithm

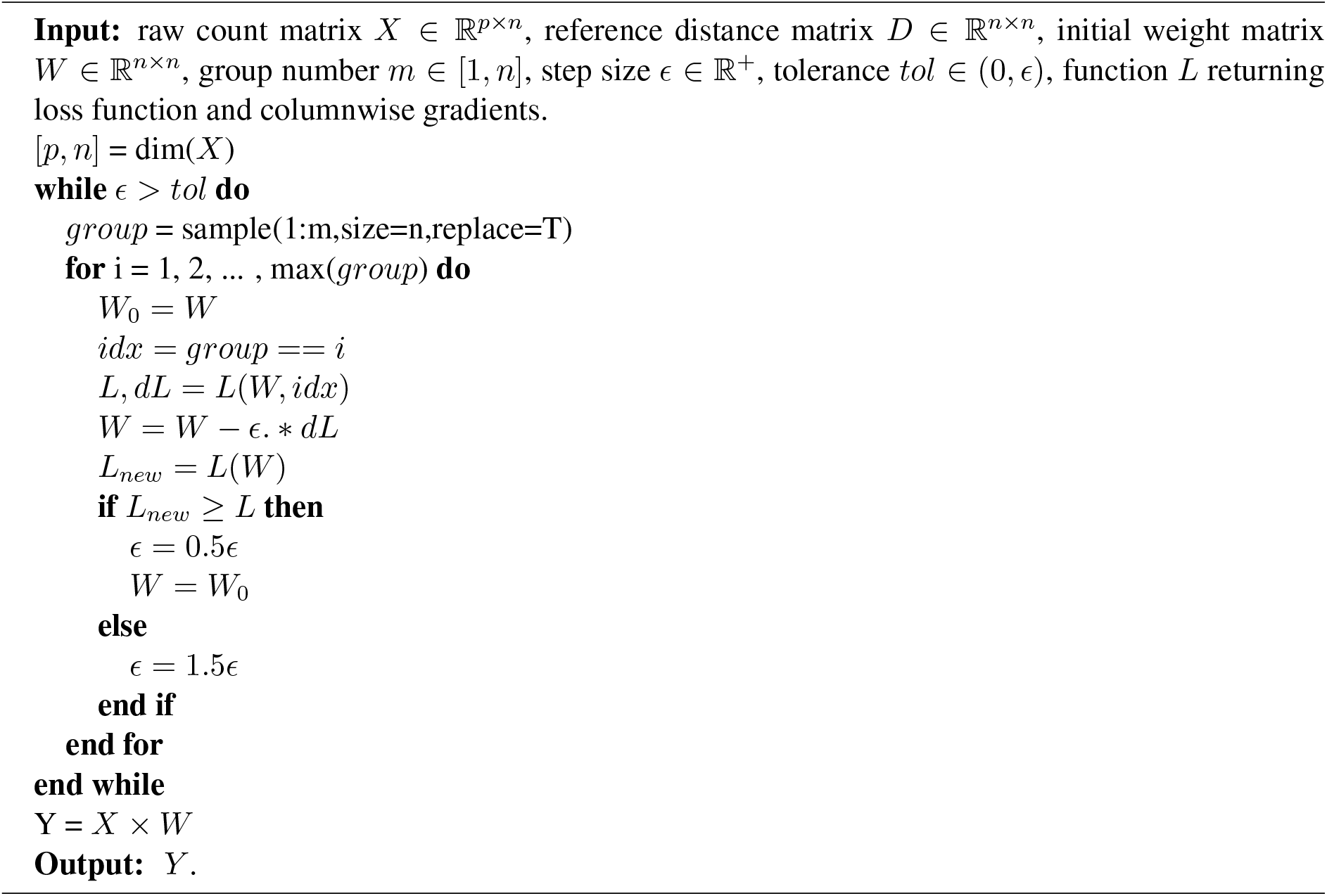

#### Random block coordinate descent algorithm

We adapt a flexible gradient descent algorithm (Algorithm 1). In each iteration, the algorithm first randomly partitions *n* subjects into *m* groups. Then gradient descent is sequentially conducted from group 1 to group *m* to update the group-specific columns in *W*. That is, the subjects are randomly partitioned in *m* group blocks in each iteration to improve the robustness of gradient descent. Note the number of groups *m* can be customized as any integer from 1 to sample size *n*. When *m* = *n*, the algorithm is equivalent to the traditional gradient descent algorithm; when *m* = 1, the algorithm is equivalent to the coordinate descent algorithm (Wright, 2015). The flexibility alleviated both the long running time of coordinate gradient descent algorithm (Wright, 2015) and the underperformed result of gradient descent algorithm. In order to dynamically adjust the learning rate, we utilized Armijo line search (Armijo, 1966). The algorithm is stopped when the step size decreases below a threshold *tol*, indicating the approximation of a local minimum.

### Simulation design

We applied Bioconductor package `splatter` (Zappia et al., 2017) to simulate single-cell RNA-seq data. Each dataset consisted of 1000 cells and 10000 genes from four biological groups. The biological groups were in a fixed proportion 4:3:2:1. The 1000 cells were randomly allocated to four batches with equal size (250 cells) with potentially different batch effect mechanisms controlled by batch location and scale parameters. The probability of a gene being differentially expressed was fixed as 0.1. The `splatter` package reported DE factors for each gene in each group, which were used to establish the gold standard for evaluation. Given two groups of interest, if the ratio of DE factors for the two groups was larger than 3/2 or smaller than 2/3, then the gene was regarded differentially expressed among the two groups.

We considered four different configurations of the location and scale parameters of batch effects in simulation studies: (I) Location was fixed as 0.1 and scale was fixed as 0.1 for all four batches. (II) Location was 0.1, 0.2, 0.05, 0.15 respectively for the four batches, while scale was fixed 0.1. (III) Location was fixed 0.1 and scale was 0.05, 0.1, 0.25, 0.3 respectively for the four batches. (IV) Location was 0.1, 0.2, 0.05, 0.15 and scale was 0.05, 0.1, 0.25, 0.3, for the four batches respectively. (V) There were no batch effects. The location parameter decided the distance between different batches, while the scale parameter controlled the shape of each batch. Under configuration (I) the batch effects followed the same mechanism for each batch, thus it was less challenging to correct. In contrast, the batch effect mechanism varied from batch to batch under configuration (IV), which increased the difficulty of batch effect correction. Moreover, configuration (V) examined if the correction methods could maintain reasonable performances when the batch effects were negligible. For each of the five configurations, we simulated 50 datasets to be corrected by the three correction methods. For MNN, default hyperparameters were used. For scBatch, the hyperparameter *m* was chosen to be 5.

### Datasets and preprocessing

#### ENCODE human and mouse tissue data

Generated by Lin et al. (2014), the data was reanalyzed by Gilad and Mizrahi-Man (2015). We utilized the same dataset used in the reanalysis, which is available at Zenodo (Mizrahi-Man and Gilad, 2015, accessed on Feb 17 2019). The data we analyzed consisted of 26 subjects and 10290 genes.

#### Mouse neuron data

The data were generated by Usoskin et al. (2015). The raw data can be obtained from the NCBI Gene Expression Omnibus (GEO) with accession number GSE59739. We utilized a processed dataset from the public data repository of Hemberg Group (https://hemberg-lab.github.io/scRNA.seq.datasets/), where normalization, outlier exclusion and log transformation were conducted to obtain a dataset with 622 cells and 25334 genes. We further removed two batches with too few samples. The final data used for batch correction consisted of 610 cells and 25334 genes.

#### Human pancreas data

The data were generated by Xin et al. (2016). The raw data is available at the NCBI Gene Expression Omnibus (GEO) with accession number GSE81608. The data used for analysis was also obtained from Hemberg Group’s repository (https://hemberg-lab.github.io/scRNA.seq.datasets/). We used the same gene filter mentioned in Xin et al. (2016) and retained genes with RPKM counts greater than 100 in no less than 10 samples. Only cells from healthy donors were selected in batch correction and downstream analysis. The processed data contained 651 cells and 6797 genes.

### Analysis and performance evaluation scheme

#### Clustering analysis

For the bulk RNA-seq data (Mizrahi-Man and Gilad, 2015, accessed on Feb 17 2019), we applied hierarchical clustering due to the small sample size. For the single-cell RNA-seq datasets, k-means clustering was repeatedly conducted. Based on the cell subtype labels provided in raw data, we utilized Adjusted Rand Index (ARI) (Hubert and Arabie, 1985) to evaluate the agreement between the provided labels and the clustering results. The ARI index equals one if the clustering result perfectly matches the cell labels, while the index values around zero under random assignment.

#### Differential expression analysis

We applied `Seurat` (Satija et al., 2015) to conduct DE gene tests and adjusted the p-values by Benjamini and Hochberg approach (Benjamini and Hochberg, 1995). In simulation studies, we base on the gold standard to directly calculate area under the receiver operating characteristic curve (AUC) and area under the precision-recall curve (PR-AUC) to compare DE detection results. For real data, Venn diagrams are generated for genes with adjusted p-values < 10^−6^ and log fold-changes > 2 to check the agreements among different count matrices. Functional analysis of the DE gene lists are conducted using the `GOstats` package (Falcon and Gentleman, 2007), which conducts tests of over-representation of gene sets using hypergeometric test.

### Code availability

We implemented the algorithm in the open-source R package `scBatch`, which is available on GitHub (https://github.com/tengfei-emory/scBatch). The code to generate results and figures in this paper is available on GitHub (https://github.com/tengfei-emory/scBatch-paper-scripts).

## Acknowledgements

This work was supported by the following funding: NIH R01GM124061. This study was conceived of and led by T.Y. Jointly with T.Y., T.F. designed the model and algorithm, implemented the scBatch software, and led the data analysis. T.F. and T.Y. wrote the paper.

