## Supplemental Material for "Batch Effect Correction of RNA-seq Data through Sample Distance Matrix Adjustment"

Teng Fei[1] and Tianwei Yu[1]

1 Department of Biostatistics and Bioinformatics, Rollins School of Public Health, Emory University.

#### Table of Contents

1 Comparisons in computing time for ComBat, MNN and scBatch ........ 2

2 Index of Supplemental Files ........ 3

### 1 Comparisons in computing time for ComBat, MNN and scBatch

We conducted further simulation study to compare the computational cost for ComBat, MNN and scBatch. The simulation was conducted on a high performance computing workstation with Intel® Xeon® Gold 6136 CPU. Specifically, for four sample size configurations: 100, 500, 1000 and 2000, we generated 50 datasets with 20,000 genes using the same strategy as configuration (I) mentioned in Methods section. Under each sample size configuration, we recorded the time needed to conduct correction using ComBat, MNN and scBatch for each of the 50 datasets. **Supplemental Fig. S1** displays recorded running time for the three algorithms.

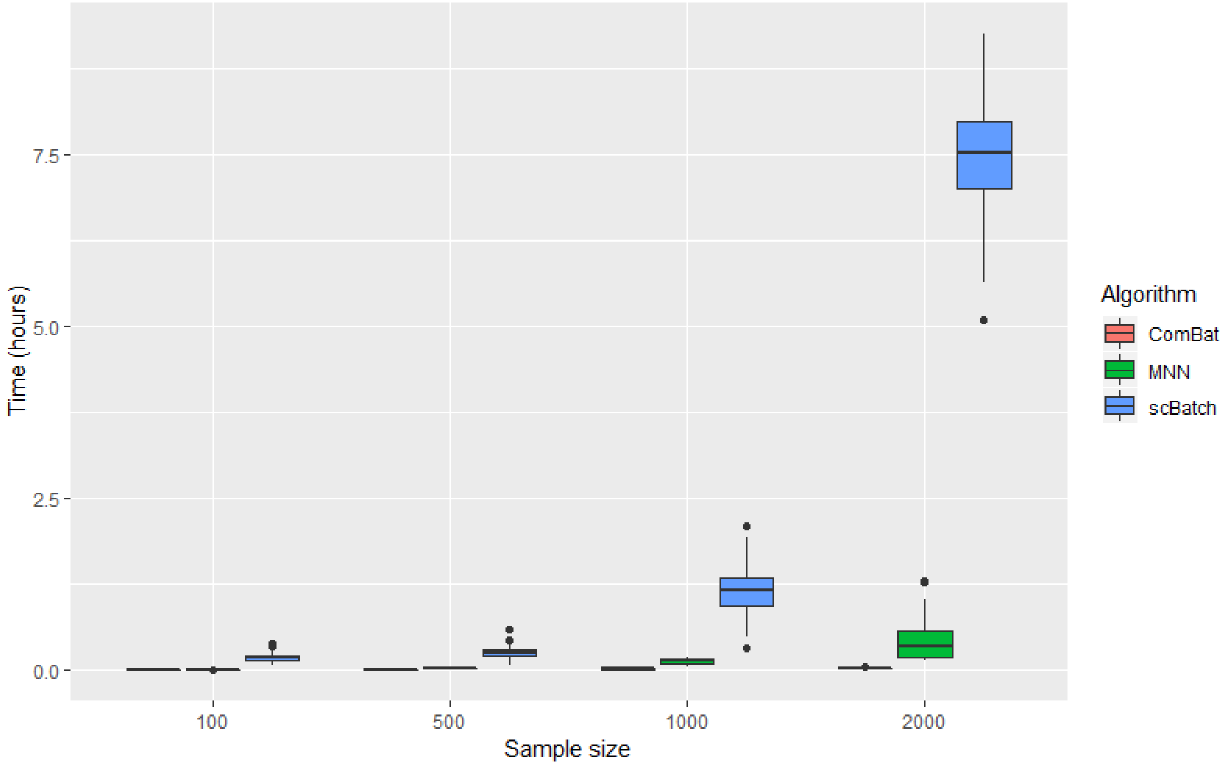


**Supplemental Fig. S1**: Boxplots of running time using ComBat, MNN and scBatch on data with different sample sizes. For each sample size configuration, 50 datasets with 20000 genes were simulated to obtain running time for the three algorithms.

#### 2 Index of Supplemental Files

**Supplemental File 1**: Full GOstats results for mouse neuron data GSE59739 (MS Excel table).

**Supplemental File 2**: Full GOstats results for human pancreas data GSE81608 (MS Excel table).

In the Excel files, each sheet contains results for one pair of cell types obtained from uncorrected data, ComBat corrected data, MNN corrected data and scBatch corrected data.
